# DecoderTCR: Compositional Pretraining and Entropy-Guided Decoding for TCR-pMHC Interactions

**DOI:** 10.64898/2026.02.04.703820

**Authors:** Boqiao Lai, Melissa Englund, Ramit Bharanikumar, Isabel Nocedal, Ali Davariashtiyani, Jason Perera, Aly A. Khan

## Abstract

Modeling recognition between T-cell receptors (TCRs) and peptide-MHC (pMHC) complexes is a fundamental challenge in computational immunology, constrained by sparse paired interaction data relative to abundant unpaired sequences. We introduce DecoderTCR, a masked language model framework that addresses this through two contributions: (1) a compositional continual pre-training curriculum that learns component representations from marginal data before refining cross-chain dependencies from limited pairs, and (2) Iterative Entropy-Guided Refinement (IEGR), a non-autoregressive decoding algorithm that resolves high-confidence positions first to provide context for uncertain regions. On held-out benchmarks, DecoderTCR achieves 0.96 AUROC for zero-shot pMHC binding prediction and 0.76 AUROC for epitope-specific TCR recognition, approaching supervised baselines without epitope-specific training. Learned representations recover structural contacts without coordinate supervision, and generated sequences exhibit realistic recombination statistics. Experimental validation reveals a prediction-generation gap: strong discrimination does not yet yield reliable generation, highlighting an open challenge for the field.

## 1. Introduction

The specificity of adaptive immunity is determined by molecular recognition between T-cell receptors (TCRs) and peptide-major histocompatibility complexes (pMHCs). This interaction underlies protective responses to infection and cancer and provides the mechanistic basis for T-cell therapies that redirect immune recognition toward disease-associated antigens. A central goal in computational immunology is to predict and design TCRs that recognize a desired pMHC target with high specificity.

Protein language models (pLMs) trained on large corpora learn structural and functional regularities from sequence alone (Lin et al., 2023). However, applying generic pLMs to immune recognition faces two challenges. First, TCR-pMHC recognition is mediated by a multi-component interface involving TCR *α/β* chains, peptide, and MHC complex. Models must capture cross-chain dependencies rather than marginal constraints within individual sequences. Second, generative design for such interfaces is intrinsically conditional and localized, requiring design of specific regions such as CDR loops while conditioning on a fixed antigenic context.

These challenges are compounded by severe data heterogeneity: TCR repertoire studies yield *∼*10^7^ TCR sequences and mass spectrometry provides *∼*10^6^ pMHC ligandome interactions, yet paired interaction data remains orders of magnitude smaller (*<* 10^5^), noisy, and biased toward common alleles and peptides (Figure 1A). This motivates a multi-stage continual pretraining framework (Ke et al., 2023; Gururangan et al., 2020) that leverages component-level representation learning from abundant marginal data before refining cross-component dependencies from limited paired interactions.

**Figure 1:**
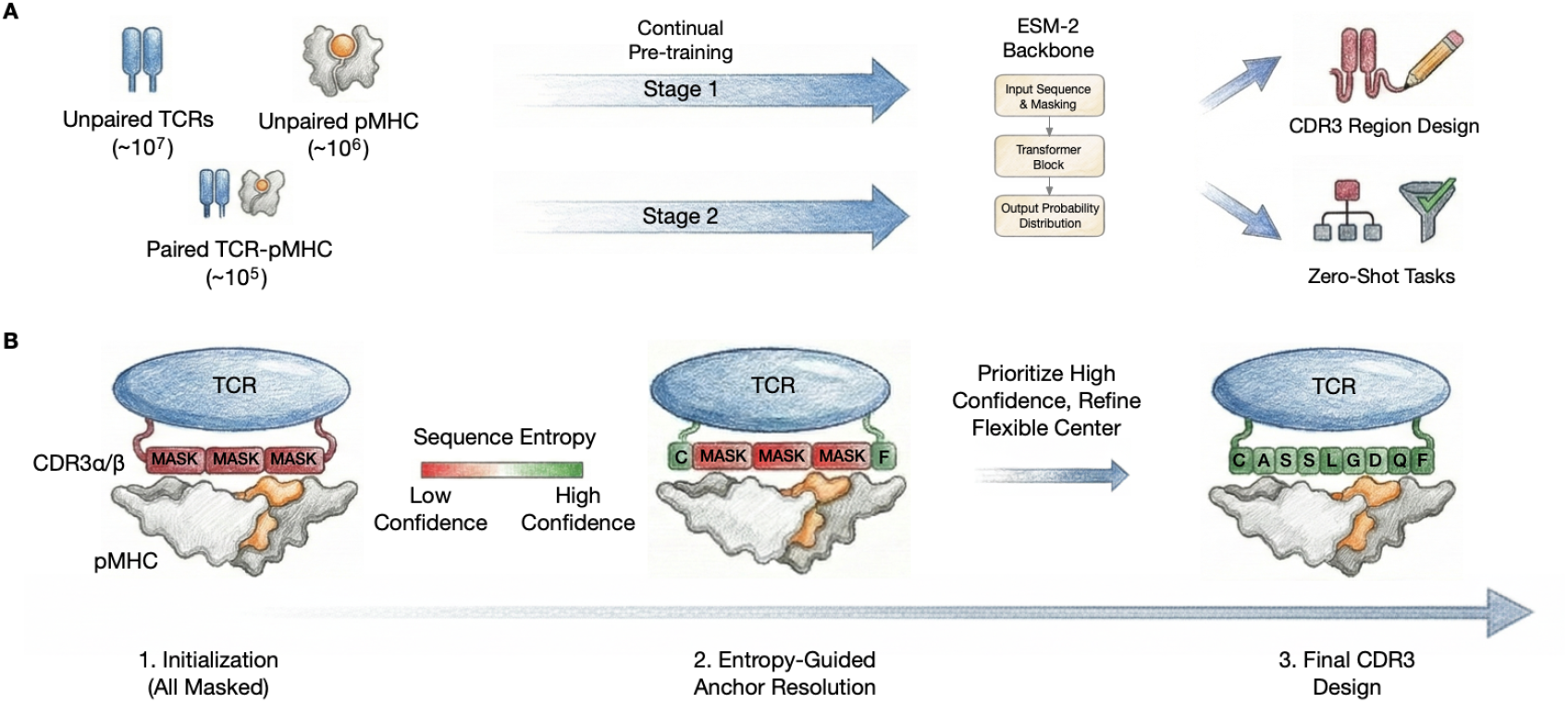
Overview of DecoderTCR. **(A)** Component-specific masked modeling integrates abundant unpaired TCR repertoires (*∼*10^7^) and pMHC data (*∼*10^6^) with sparse paired TCR-pMHC interactions (*∼* 10^5^) via tailored masking rates. **(B)** Iterative Entropy-Guided Refinement (IEGR) performs constrained CDR3 redesign by resolving low-entropy anchor positions first, followed by local resampling of flexible regions.

Prior discriminative methods learn binding classifiers (Montemurro et al., 2021; Lu et al., 2021; Weber et al., 2021) but cannot directly generate sequences. Autoregressive protein language models (Madani et al., 2023; Ferruz et al., 2022) can sample novel sequences, but left-to-right factorization is poorly suited for constrained loop redesign where residues are jointly constrained by both N-terminal and C-terminal context. Structure-conditioned methods (Dauparas et al., 2022; Watson et al., 2023) require coordinate supervision which remains scarce for TCR-pMHC interfaces (< 10^3^ solved structures versus *>* 10^4^ paired sequences).

We introduce **DecoderTCR**, a sequence-first framework addressing TCR-pMHC *prediction* and *design* through two innovations (Figure 1). First, we propose a **multi-stage continual pre-training curriculum** that bridges heterogeneous data sources. Rather than learning the joint distribution *p*(,) from sparse paired data alone, we continually pretrain ESM-2 with component-specific masking schedules tailored to each immune component. This strategy allows abundant marginal data to regularize learning while limited paired data refines cross-chain dependencies.

Second, we propose **Iterative Entropy-Guided Refinement (IEGR)**, a non-autoregressive decoding strategy for constrained CDR3 redesign. Building on confidence-based iterative decoding (Chang et al., 2022; Ghazvininejad et al., 2019), IEGR iteratively unmasks positions in order of predictive entropy, resolving low-entropy anchor residues first to establish a stable scaffold, then refining high-entropy flexible regions through local masked resampling. This schedule respects the biophysical hierarchy where conserved positions constrain hypervariable loops.

We evaluate DecoderTCR across representation quality, zeroshot binding prediction, and experimental validation. On held-out benchmarks with stringent allele and epitope splits, DecoderTCR achieves 0.96 AUROC on zero-shot pMHC binding (compared to 0.45 for ESM-2) and 0.76 AUROC on epitope-specific TCR recognition, approaching supervised baselines trained explicitly on target epitopes. Peptide interaction scores localize structural contact positions despite training without coordinate supervision. Experimental screening of designed TCRs confirms that zero-shot generation remains challenging, highlighting the gap between strong prediction and reliable generation.

Our contributions are:

1. **Compositional continual pre-training**. A two-stage curriculum bridging the 100*×* gap between marginal sequences (*∼*10^7^ TCRs, *∼*10^6^ pMHCs) and paired interactions (*∼*10^5^) via functionally-informed masking and implicit experience replay. To our knowledge, this yields the first multi-component protein language model capturing the full immune synapse.
2. **Entropy-guided decoding with span scoring**. Iterative Entropy-Guided Refinement (IEGR) resolves low-entropy anchors before high-entropy hypervariable regions, producing sequences with realistic recombination statistics and structural plausibility. Span pseudo-likelihood scoring enables binding prediction and recovery of structural contacts without coordinate supervision.
3. **Comprehensive evaluation**. Zero-shot generalization on strictly held-out alleles and epitopes, wet-lab validation of designed TCRs, and rigorous characterization of the prediction-generation gap.

## 2. Related Work

### Protein Language Models for Immune Repertoires

Large-scale protein language models demonstrate that structural and functional constraints can be learned from sequence data (Lin et al., 2023; Elnaggar et al., 2021). Immune-focused models including TCR-BERT (Wu et al., 2024), TCRLM (Fang et al., 2024), ESMCBA (Mares et al., 2025), and IgLM (Shuai et al., 2023) encode priors tailored to adaptive immune receptors, supporting repertoire-level tasks such as clonotype clustering and motif discovery. A key limitation is that standard pretraining captures marginal regularities within single sequence families rather than conditional dependencies across multi-component interfaces. Our component-specific masking strategy combines marginal priors from abundant unpaired data with cross-chain dependencies from limited paired interactions.

### TCR-pMHC Binding Prediction

Discriminative methods have advanced from hand-crafted similarity kernels (Dash et al., 2017) to deep learning approaches including NetTCR-2.0 (Montemurro et al., 2021), ERGO-II (Springer et al., 2021), pMTnet (Lu et al., 2021), TITAN (Weber et al., 2021), and ATM-TCR (Cai et al., 2022). While effective on seen epitopes, these methods degrade under distribution shift to unseen epitopes or rare alleles (Lu et al., 2025) and cannot directly generate sequences. Structure-conditioned methods (Dauparas et al., 2022; Watson et al., 2023) require coordinate supervision scarce for TCR-pMHC interfaces. DecoderTCR enables generative sampling while leveraging unpaired data to improve generalization.

### Continual Pre-training

Continual pre-training adapts foundation models to specialized domains (Gururangan et al., 2020). Curriculum strategies can mitigate catastrophic forgetting when adapting to new distributions (Ke et al., 2023). Domain-adaptive pre-training on scientific (Beltagy et al., 2019) and biomedical (Lee et al., 2020) text improves downstream performance. Our two-stage curriculum extends this paradigm to multi-component protein interfaces where heterogeneous data sources leverage distinct masking strategies.

### Non-Autoregressive Decoding

Non-autoregressive generation mitigates the sequential bottleneck of autoregressive models. Mask-Predict (Ghazvininejad et al., 2019) introduced iterative masked decoding for translation. MaskGIT (Chang et al., 2022) proposed confidence-based token selection for iterative parallel decoding. IEGR adapts this approach to constrained protein design with entropy-guided position selection and block-wise Gibbs refinement.

## 3. Methods

We formulate TCR-pMHC modeling as learning conditional distributions over multi-component immune complexes. Our approach comprises four components: a unified sequence representation, component-aware masked language modeling, span pseudo-likelihood scoring for binding prediction and interpretability, and entropy-guided refinement for generation.

### Notation

Let denote the amino acid vocabulary with ‖ = 20 canonical residues. We denote pMHC sequences as = (*x*_1_, …, *x*_*n*_) *∈*^*n*^ and TCR sequences as = (*y*_1_, …, *y*_*m*_) *∈*^*m*^. The joint TCR-pMHC complex is represented as concatenation *⊆*= [∥] *∈*^*n*+*m*^. For sequence = (*s*_1_, …, *s*_*L*_) and position set ⊆ [*L*]:= {1, …, *L*}, we write for the sequence with all positions in replaced by.

### 3.1. Unified Multi-Chain Representation

To capture cross-chain dependencies, we represent the TCRpMHC complex as a single concatenated sequence:

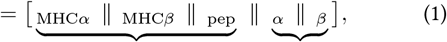

where _*α*_,_*β*_ are the TCR *α* and *β* chains, _pep_ is the presented peptide, and _MHC_ comprises MHC chains (heavy chain and *β*_2_-microglobulin for Class I, or *α/β* chains for Class II).

### 3.2. Two-Stage Continual Pre-training Curriculum

We adopt a continual pre-training strategy organized as a two-stage curriculum, progressing from large-scale pMHC and TCR sequence pre-training to paired TCR–pMHC interaction pre-training. Rather than uniform masking, we use functionally informed stage specific masking schedule that concentrates self-supervision on binding-relevant positions.

#### Stage 1: Component-specific representations

We train jointly on TCR repertoires from OTS (Raybould et al., 2024) and pMHC ligandomes from MHC Motif Atlas (Tadros et al., 2023).

The Stage 1 objective is:

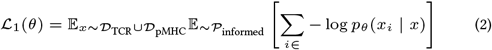

where 𝒫_informed_ is a position-dependent masking distribution based on component specific functional annotations.

#### Stage 2: Cross-chain interaction learning

We continue training on paired TCR-pMHC sequences from VDJdb (Bagaev et al., 2020). The key insight is that paired data allows the model to learn how CDR sequences and peptide sequences constrain each other through the binding interface. We use joint masking of CDR and peptide positions:

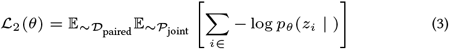

where *𝒫*_joint_ is a joint masking distribution spanning all components. See Appendix A.1 for masking details and Appendix A.2 for optimization configuration.

This Stage 2 objective subsumes Stage 1 tasks. When peptide positions are masked, the model predicts them from MHC context. When CDR positions are masked, the model predicts them from framework context. The additional signal comes from cross-component conditioning, where CDR predictions leverage peptide context and vice versa. This compositional structure provides implicit experience replay, maintaining Stage 1 competencies while learning interaction-specific features.

### 3.3. Span Pseudo-Log-Likelihood Scoring

Standard pseudo-log-likelihood (PLL) masks single positions independently. For binding prediction, we instead use *span* PLL (sPLL), which masks entire functional regions simultaneously (Joshi et al., 2020). This better captures that binding interfaces function as coherent units. We derive two uses of sPLL: *aggregate scores* for binding prediction and *per-position scores* for structural interpretation.

#### Aggregate sPLL for binding prediction

To predict binding compatibility, we mask all positions in a functional span and average their log-probabilities. Let _pep_ denote the set of peptide position indices. For pMHC binding:

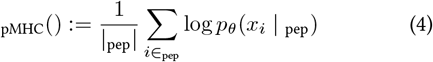

where _pep_ denotes the pMHC sequence with peptide positions replaced by and MHC positions unchanged. Following Goyal et al. (2021), higher sPLL indicates sequences more compatible with the learned distribution, though sPLL does not correspond to a normalized probability.

For TCR-pMHC recognition, we include TCR context:

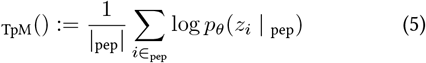

where peptide positions _pep_ are indexed identically in both and = [∥].

#### Per-position sPLL for structural interpretation

To identify *which residues* drive predictions, we decompose sPLL into per-position contributions inspired by pointwise mutual information (PMI). For consistent distributions, PMI(*x*; *y*) = log *p*(*x*|*y*)*−* log *p*(*x*) quantifies information gain from conditioning. Since MLMs do not define consistent joints (Goyal et al., 2021), we use a pseudo-likelihood analogue comparing predictions under different contexts.

#### Intuition for pMHC binding

The pMHC interaction score measures whether MHC context improves peptide prediction over background. High scores indicate positions where the MHC groove constrains amino acid identity, which are typically anchor positions P2 and PΩ (C-terminus). For HLA-A*02:01, these anchors are characteristically leucine at P2 and valine at PΩ.

#### Definition 3.1 (pMHC Interaction Score). For peptide position *i ∈*_pep_

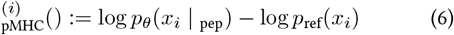

where *p*_ref_ is the uniform distribution over the 20 standard amino acids

#### Intuition for TCR recognition

The TCR-pMHC interaction score measures additional predictive information from TCR presence beyond MHC alone. High scores indicate positions where TCR directly contacts the peptide, which are typically solvent-exposed central positions P4–P7 that engage CDR3 loops.

#### Definition 3.2 (TCR-pMHC Interaction Score). For peptide position *i ∈*_pep_

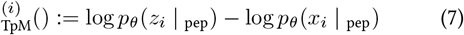

Positive scores indicate positions where TCR context improves peptide prediction.

### 3.4. Inference: Iterative Entropy-Guided Refinement

Given a target pMHC and a TCR scaffold (fixed V/J genes and framework regions), we aim to design CDR3*α* and CDR3*β* sequences that jointly confer specific recognition. This is a constrained generation problem where the designed regions must be compatible with both scaffold context and target antigen.

Standard approaches have limitations. Autoregressive generation imposes arbitrary left-to-right ordering that ignores bidirectional constraints from framework regions. Single-pass parallel decoding ignores dependencies between designed positions. We propose **Iterative Entropy-Guided Refinement (IEGR)**, adapting confidence-based non-autoregressive decoding (Chang et al., 2022; Ghazvininejad et al., 2019) to constrained protein design. The key insight is that resolving high-confidence positions first provides reliable context for subsequent predictions.

#### Biological motivation for entropy ordering

Both CDR3*α* and CDR3*β* loops contribute to the TCR-pMHC interface, with CDR3*β* typically making dominant peptide contacts while CDR3*α* modulates specificity. Designing both loops jointly ensures interface compatibility. CDR3 loops share conserved boundary residues, including N-terminal cysteines and C-terminal Phe-Gly or Trp-Gly motifs, that anchor loop geometry. Central positions are hypervariable and determine antigen specificity.

Entropy-guided ordering naturally respects this hierarchy. Anchor positions have low predictive entropy and are resolved first, establishing a stable scaffold. High-entropy central positions are resolved last, conditioned on established anchors. By computing entropy across both loops simultaneously, the algorithm interleaves decoding between chains, selecting whichever position has lowest uncertainty at each step. This allows cross-chain dependencies to inform the design of both loops.

#### Algorithm

Let _0_ denote design positions (CDR3 residues). For masked positions _*t*_ at step *t*, we quantify uncertainty via predictive entropy:

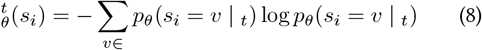

#### Phase 1: Entropy-Guided Construction

Starting from all design positions masked, we iteratively select the position with lowest entropy (highest confidence), sample its value from the predicted distribution with temperature *τ*, and remove it from the mask set. This continues until all positions are filled.

##### Algorithm 1

Iterative Entropy-Guided Refinement (IEGR)

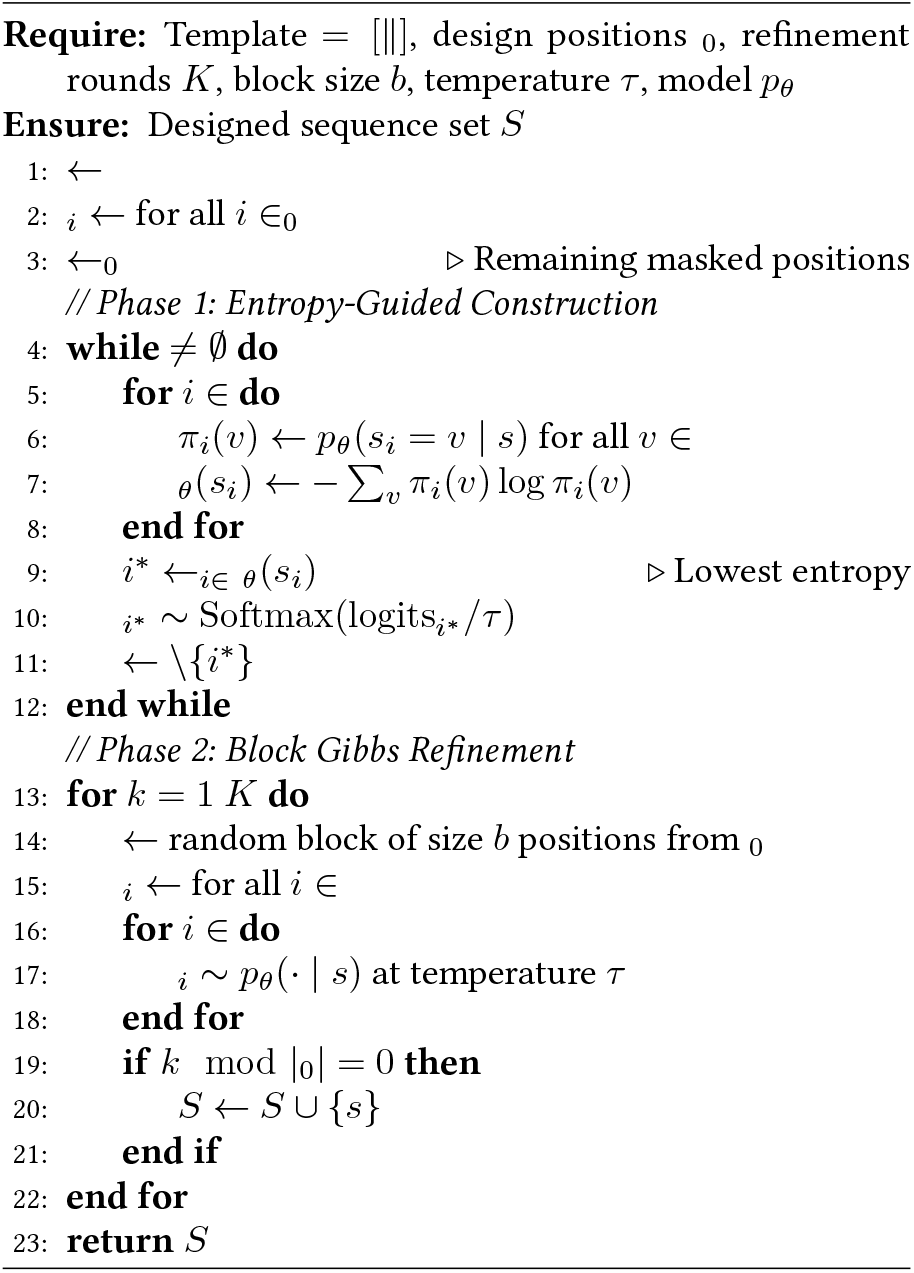

#### Phase 2: Block Gibbs Refinement

After initial construction, we perform *K* rounds of local refinement. Each round selects a random block of *b* positions, masks them, and resamples. This enables escape from suboptimal configurations.

Since masked language model conditionals need not arise from a consistent joint distribution (Goyal et al., 2021), IEGR lacks classical MCMC guarantees. However, the entropy-guided ordering can be interpreted as a greedy approximation to maximizing mutual information between resolved and unresolved positions at each step. We view IEGR as a practical search heuristic whose effectiveness we evaluate empirically. Full pseudocode appears in Algorithm 1; implementation details and hyperparameters are provided in Appendix B.

## 4. Experimental Setup

### Training Data

Stage 1 trains on TCR repertoires (1.5 *×*10^6^ paired TCR *αβ* sequences from OTS (Raybould et al., 2024)) and pMHC ligandomes (1.0*×* 10^6^ peptide-MHC pairs from two large monoallelic cell line datasets, and the MHC Motif Atlas (Tadros et al., 2023; Sarkizova et al., 2020; Abelin et al., 2017) combined with synthetic peptides consisting of high-confidence predictions from MixMHCpred-2.2 (Gfeller et al., 2023). Stage 2 trains on paired TCR-pMHC interactions (3.0 *×*10^4^ from VDJdb (Bagaev et al., 2020)). Optimization hyperparameters and computational cost are detailed in Appendix A.

### Evaluation Splits

We use strict held-out splits to evaluate generalization. For pMHC binding, we held out 10 Class I alleles and 6 Class II alleles completely from Stage 1. See Table 6 for a complete list. For TCR-pMHC recognition, we held out epitopes YLQPRTFLL (SARS-CoV-2 Spike) and GLCTLVAML (EBV BMLF1) and all TCRs recognizing these epitopes from Stage 2. Full details in Appendix C.

### Models and Baselines

DecoderTCR uses ESM-2 backbones at 650M and 3B parameters. For pMHC binding, we compare against MHCflurry 2.0 (O’Donnell et al., 2020), MixMHCpred-2.2 (Gfeller et al., 2023), NetMHCpan4.1 (Reynisson et al., 2020). Zero-shot ESM scoring is performed similarly as 4. For TCR-pMHC recognition, we compare against NetTCR-2.2 (Jensen and Nielsen, 2023). For generation evaluation, we use AlphaFold3 (Abramson et al., 2024) interface pTM (ipTM) as an in silico oracle.

## 5. Results

We evaluate DecoderTCR on representation quality, knowledge retention, binding prediction, structural interpretability, and experimentally validated de novo design.

### 5.1. pMHC Binding Prediction

Following Stage 1 training on joint pMHC ligandomes and TCR repertoires with informed masking, we assess model quality along two dimensions: (1) zero-shot generalization to unseen HLA alleles and (2) representation transferability for supervised pMHC binding prediction, both prior to Stage 2 training.

#### Zero-shot generalization

We rank peptides by aggregate sPLL (Eq. 4). Table 1 reports performance on 16 HLA alleles excluded from training. ESM-2, despite training on billions of protein sequences, performs near random (AUROC 0.41-0.56), confirming that generic protein representations do not encode MHC-specific binding preferences. Stage 1 DecoderTCR achieves AUROC 0.93-0.96 on Class I and 0.90-0.91 on Class II, demonstrating that domain-specific pretraining yields strong zero-shot transfer. The modest Class I versus Class II gap reflects biological differences in binding groove conservation (Jones et al., 2006). Per-allele ROC and PR curves are shown in Appendix Figure 5.

**Table 1:**
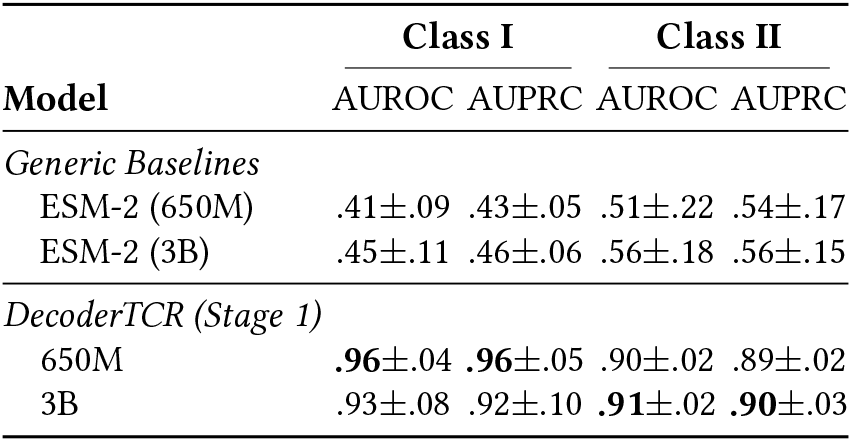
Zero-shot pMHC binding. Stage 1 trains jointly on pMHC and TCR data. Test alleles completely excluded from training. Mean ± SD across 10 Class I and 6 Class II alleles.

| Model | Class I |  | Class II |  |
| --- | --- | --- | --- | --- |
|  | AUROC | AUPRC | AUROC | AUPRC |
| *Generic Baselines* |  |  |  |  |
| ESM-2 (650M) | .41±.09 | .43±.05 | .51±.22 | .54±.17 |
| ESM-2 (3B) | .45±.11 | .46±.06 | .56±.18 | .56±.15 |
| *DecoderTCR (Stage 1)* |  |  |  |  |
| 650M | .96±.04 | .96±.05 | .90±.02 | .89±.02 |
| 3B | .93±.08 | .92±.10 | .91±.02 | .90±.03 |

#### Transfer to supervised prediction

To verify that representations encode transferable features, we train an MLP classifier on frozen embeddings using IEDB *pIC*_50_ binding affinity data and evaluate on ESCAPE-seq (Shi et al., 2025), an independent prospective benchmark. DecoderTCR-3B achieves AUROC 0.77, matching specialized predictors (MixMHCpred, NetMHCpan, MHCflurry) that were purpose-built with allelespecific architectures. ESM-2 achieves only 0.62-0.63 regardless of scale, confirming that model size alone does not substitute for domain-relevant pretraining (Table 2).

**Table 2:** Supervised Oncogene mutant pMHC binding prediction. MLP classifier on frozen Stage 1 embeddings, tested on prospective data from Shi et al. (2025).

| Model | AUROC | AUPRC |
| --- | --- | --- |
| *Specialized Predictors* |  |  |
| MixMHCpred-3.0 | .77±.07 | .44±.15 |
| NetMHCpan-4.1 | .77±.08 | .44±.15 |
| MHCflurry 2.0 | .76±.08 | .45±.14 |
| *ESM Embeddings* |  |  |
| ESM-2 (650M) | .62±.06 | .19±.12 |
| ESM-2 (3B) | .63±.07 | .19±.12 |
| *DecoderTCR Embeddings (Stage 1)* |  |  |
| 650M | .75±.07 | .35±.15 |
| 3B | .77±.07 | .35±.16 |

### 5.2. Knowledge Retention During Continual Learning

A central question for our two-stage continual pre-training curriculum is whether Stage 2 training on paired data overwrites Stage 1 representations. We monitor two sentinel metrics throughout Stage 2 training progression: zero-shot pMHC binding AUROC on held-out alleles and TCR inverse perplexity on held-out TCR-pMHC sequences.

Figure 2 demonstrates both metrics remain stable or improve throughout 3500 steps of Stage 2 training. Zero-shot pMHC AUROC remains constant at 0.925-0.93 with no discernible degradation. TCR inverse perplexity increases from 0.72 to 0.79 (+10%), suggesting that exposure to binding-relevant CDR sequences during Stage 2 refines rather than overwrites TCR representations. We posit this stability arises from *implicit experience replay*: each paired TCR-pMHC example contains valid pMHC and TCR subsequences, so the joint masking objective (Eq. 3) subsumes Stage 1 tasks while adding cross-component signal.

**Figure 2:**
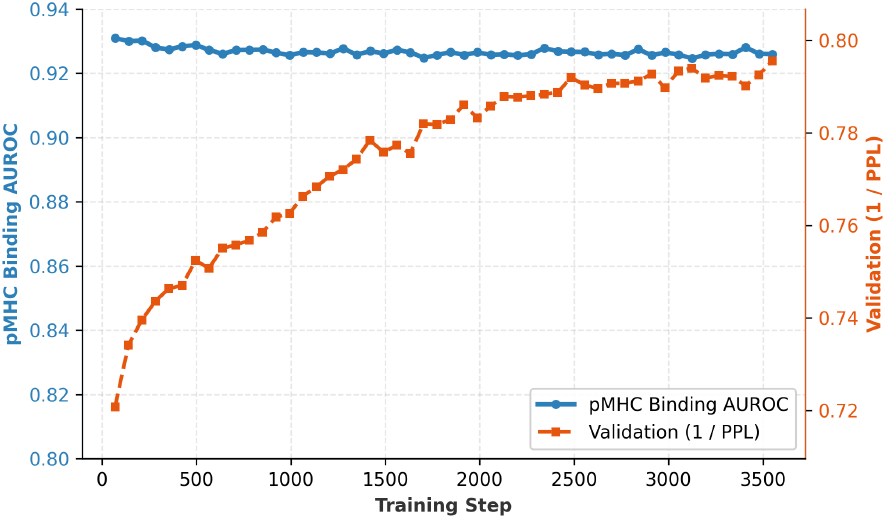
Knowledge retention during Stage 2. Zero-shot pMHC AUROC (blue) measures binding prediction on alleles excluded from all training. TCR inverse perplexity (orange) measures language modeling quality on held-out sequences.

### 5.3. Epitope-Specific TCR Recognition

We evaluate the core capability: given a fixed pMHC target, can DecoderTCR discriminate binding from non-binding TCRs? This epitope-specific setting mirrors therapeutic TCR design where the target antigen is known.

#### Evaluation protocol

We use YLQPRTFLL (YLQ:SARSCoV-2 Spike) and GLCTLVAML (GLC:EBV BMLF1), both HLA-A*02:01-restricted, with experimentally validated binders and non-binders from Messemaker et al. (2025). We prioritize these epitopes for ground-truth reliability over breadth, given that most public TCR-pMHC labels remain noisy. TCR-pMHC pairs are scored by summed sPLL over masked peptides under the joint context; higher scores indicate higher predicted binding probability. Both epitopes are completely excluded from all DecoderTCR training, enabling true zero-shot evaluation. We compare against NetTCR-2.2, a supervised baseline trained explicitly on these epitopes.

#### Both stages are necessary

Table 3 reveals that neither stage alone suffices. Stage 1 alone achieves only 0.18 AUROC because pMHC and TCR representations occupy disjoint embedding regions without paired data to align them. Stage 2 alone achieves 0.56 AUROC, functional but 0.20 below the full pipeline, indicating that Stage 1 provides beneficial initialization. The full two-stage pipeline achieves 0.76 AUROC, approaching NetTCR-2.2 (0.88) despite never seeing YLQPRTFLL during training while NetTCR-2.2 was trained explicitly on this epitope. Extended comparisons against additional supervised methods trained prior to this epitope’s characterization appear in Appendix Figure 6.

**Table 3:** Epitope-specific TCR recognition. Given a fixed pMHC, discriminate binding from non-binding TCRs. Both epitopes completely excluded from DecoderTCR training.

| Model | YLQ |  | GLC |  |
| --- | --- | --- | --- | --- |
|  | AUC | PRC | AUC | PRC |
| *Supervised (Trained on Epitope)* |  |  |  |  |
| NetTCR-2.2 | .88 | .82 | .69 | .87 |
| *Zero-Shot Baselines* |  |  |  |  |
| ESM-2 (3B) | .36 | .34 | .48 | .71 |
| *DecoderTCR (3B) Ablations* |  |  |  |  |
| Stage 1 only | .18 | .29 | .43 | .70 |
| Stage 2 only | .56 | .42 | .46 | .70 |
| Stage 1 & 2 | **.76** | **.70** | **.64** | **.83** |

**Table 4.**
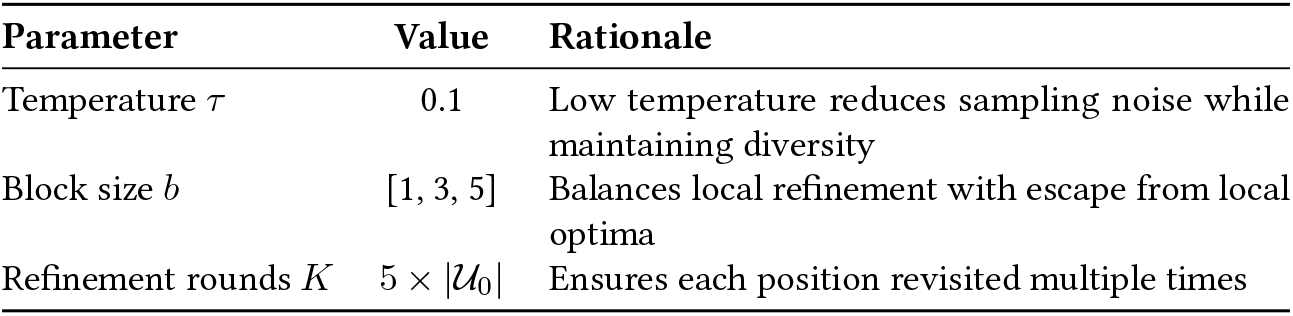
IEGR hyperparameters.

#### Ablation summary

We consolidate ablation findings. For training, both stages are necessary: Stage 1 alone fails (0.18 AUROC) because representations are not aligned; Stage 2 alone underperforms (0.56 AUROC) without regularization from unpaired data; the full pipeline Stage 1+2 achieves 0.76 AUROC. For inference, entropy-guided ordering outperforms single-pass decoding (Section 5.5). Extended ablations on masking strategies appear in Appendix D.

### 5.4. structural Interpretability via Interaction Scores

While aggregate sPLL enables binding prediction, it does not reveal *which residues* drive the prediction. We apply perposition interaction scores (Eqs. 6–7) to the held-out validated YLQPRTFLL positive binders and overlay the summarized interaction scores over a TCR-pMHC complex with solved structure (PDB 7RTR), using no structural supervision during training. By treating the peptide as the shared interface, we decompose binding signal into MHC-driven anchoring versus TCR-driven recognition.

#### TCR-Peptide interface

_TpM_ (Eq. 7) quantifies how TCR context improves peptide prediction beyond MHC alone. Positions P4-P7 show elevated scores, corresponding to the peptide bulge contacting CDR3 loops (Figure 3B). Anchor positions P2 and P9 show minimal TCR-dependent signal, as expected for residues facing the MHC groove rather than TCR.

**Figure 3:**
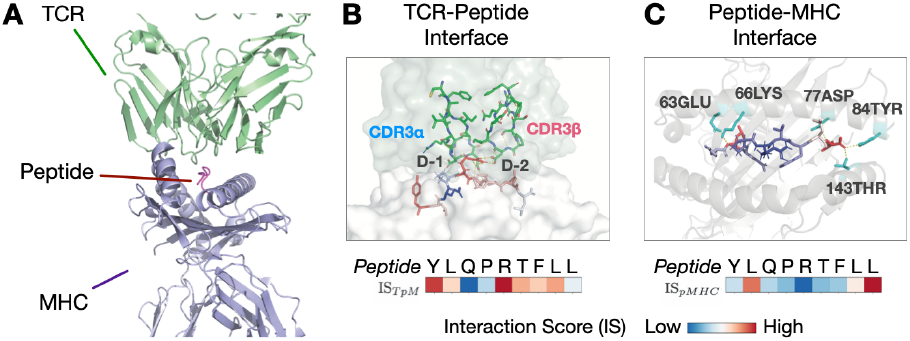
Interaction scores recover structural contacts without supervision. (A) Overview of TCR-pMHC ternary complex (PDB 7RTR) showing TCR (green) engaging peptide (pink) presented by MHC (purple). (B) TCR-peptide interface with two salt bridges between TCR*α* Asp109/TCR*β* Asp110 and peptide Arg5. TpM highlights TCR contact positions P4-P7. (C) Peptide-MHC interface with anchor residues P2 (Leu) and P9 (Leu) buried in MHC pockets. pMHC highlights anchors P2 and P9.

#### Peptide-MHC interface

_pMHC_ (Eq. 6) quantifies how MHC context improves prediction at each peptide position. Positions P2 and P9 show elevated scores, corresponding to canonical anchor residues buried in MHC binding pockets (Figure 3C). These anchors (typically leucine at P2 and valine at P9 for HLA-A*02:01) drive stable peptide presentation (Madden et al., 1993). Central positions P4-P7 show lower MHC-dependent signal.

The complementary pattern (MHC anchors at termini, TCR contacts at center) emerges purely from sequence-level training, suggesting representations capture biologically relevant interface features.

### 5.5. Generative Design and Experimental Validation

Moving from binding prediction to sequence design, we evaluate the model’s ability to zero-shot generate novel binders for a strictly held-out target. Given YLQPRTFLL/HLA-A*02:01 and a TCR scaffold (fixed V/J genes and framework regions taken from validated positive binding TCRs in (Messemaker et al., 2025)), the task is to design CDR3*α* and CDR3*β* sequences that confer target recognition in a zero-shot setting. We compare three decoding strategies: greedy (argmax), oneshot (simultaneous sampling), and IEGR (entropy-guided with Gibbs refinement).

#### In silico evaluation

Figure 4 compares strategies using AlphaFold3 ipTM as a structural proxy. Single-pass methods (greedy, oneshot) produce ipTM distributions comparable to validated non-binders. IEGR significantly outperforms both (Mann-Whitney *p <* 0.001) by resolving low-entropy anchors before high-entropy positions, enabling escape from suboptimal configurations. Sequence-level evaluation with OLGA (Sethna et al., 2019) confirms IEGR produces the highest proportion of biologically plausible sequences (Appendix Figure 7).

**Figure 4:**
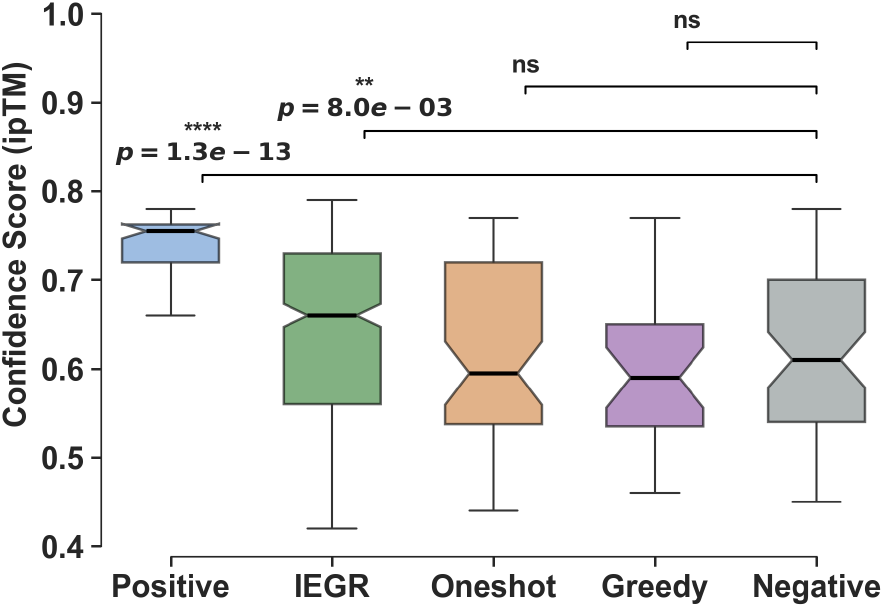
*In silico* assessment of CDR design quality. Distributions of AlphaFold3 ipTM scores for designs targeting YLQPRTFLL/HLA-A*02:01. Positive and Negative controls are taken from (Messemaker et al., 2025) validated YLQ binding TCRs. Brackets denote two-sided Wilcoxon rank-sum tests against validated negative controls. IEGR significantly outperforms non-binders (*p <* 0.001).

**Figure 5:**
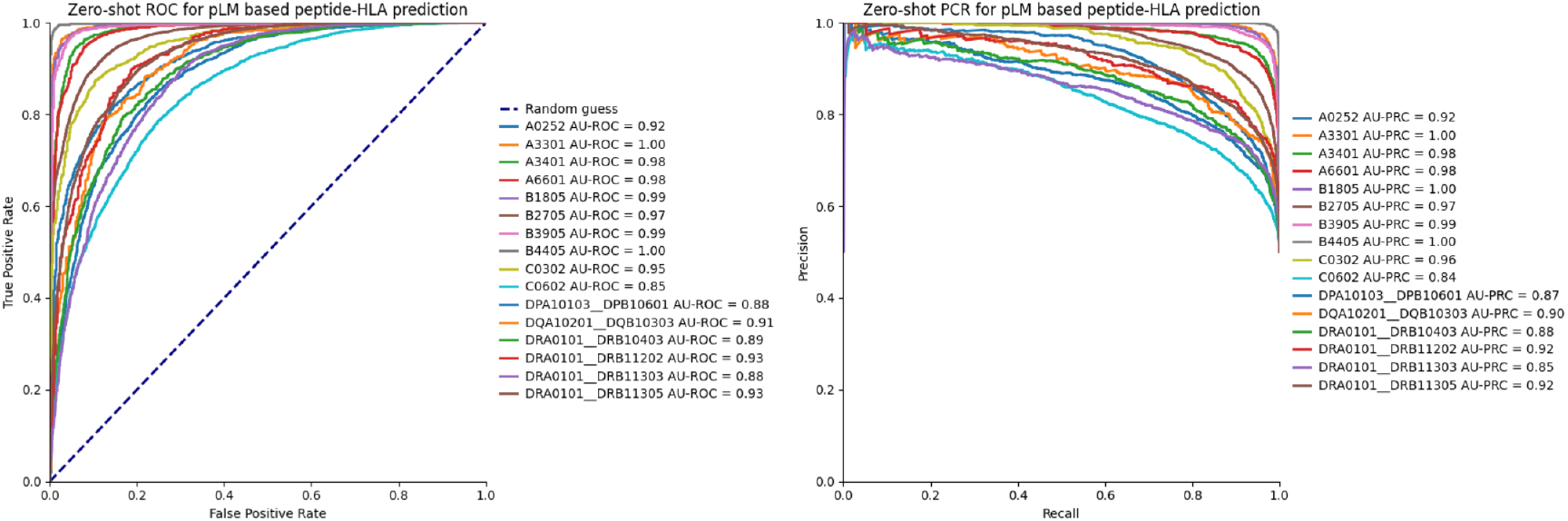
Zero-shot pMHC binding predictions by allele. Per-allele ROC (left) and PR (right) curves for Stage 1 DecoderTCR model on held-out HLA alleles. All 10 Class I alleles and 6 Class II alleles achieve AUROC ≥ 0.85, demonstrating consistent zero-shot generalization across diverse HLA types. Table 1 reports aggregate statistics.

**Figure 6:**
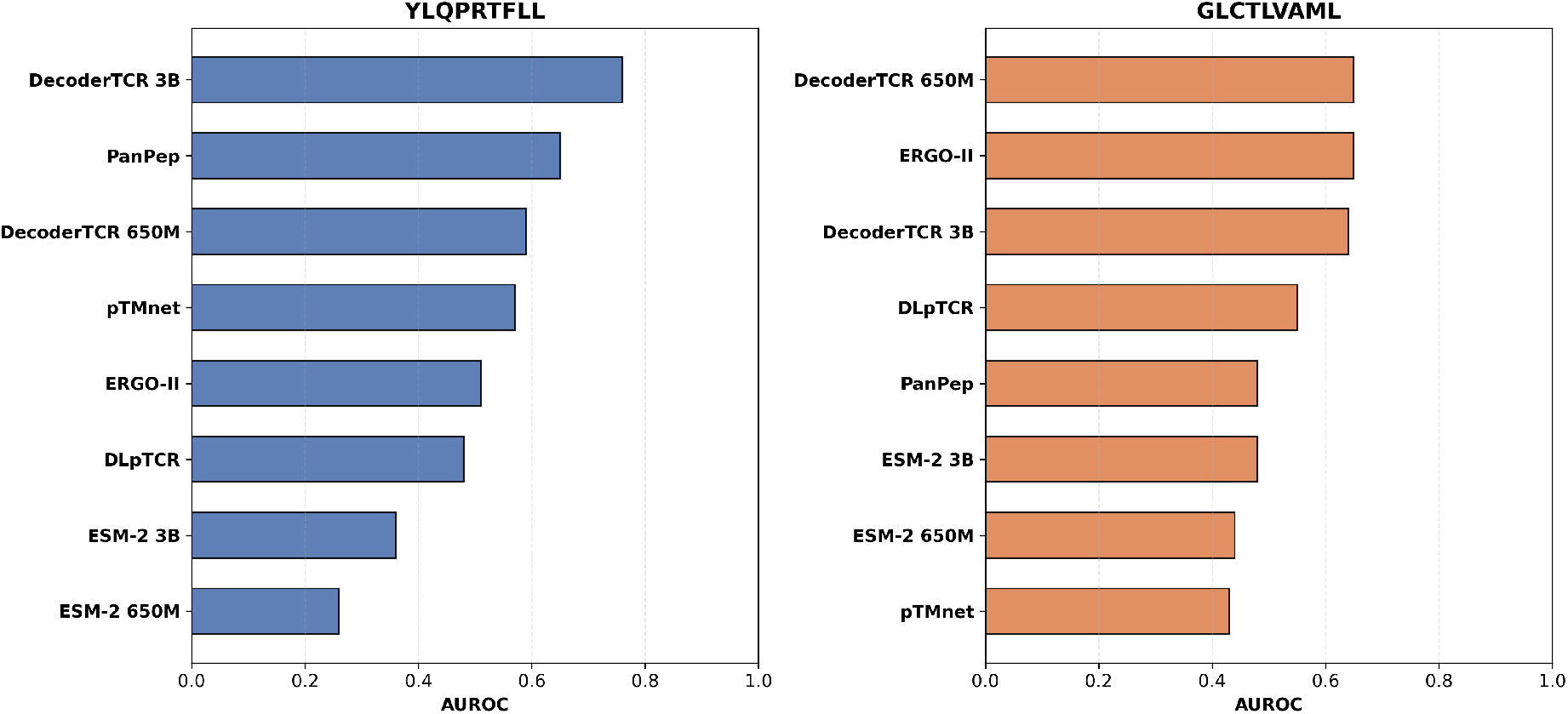
Extended TCR-pMHC recognition comparison. AUROC for discriminating binding from non-binding TCRs on YLQPRTFLL (left) and GLCTLVAML (right) epitopes. DecoderTCR is compared against other supervised methods (ERGO-II, pMTnet, DLpTCR, PanPep) and generic protein language models (ESM-2 650M/3B). Despite zero-shot evaluation, DecoderTCR approaches or exceeds supervised baselines on both epitopes. Validation data from Messemaker et al. (2025).

**Figure 7:**
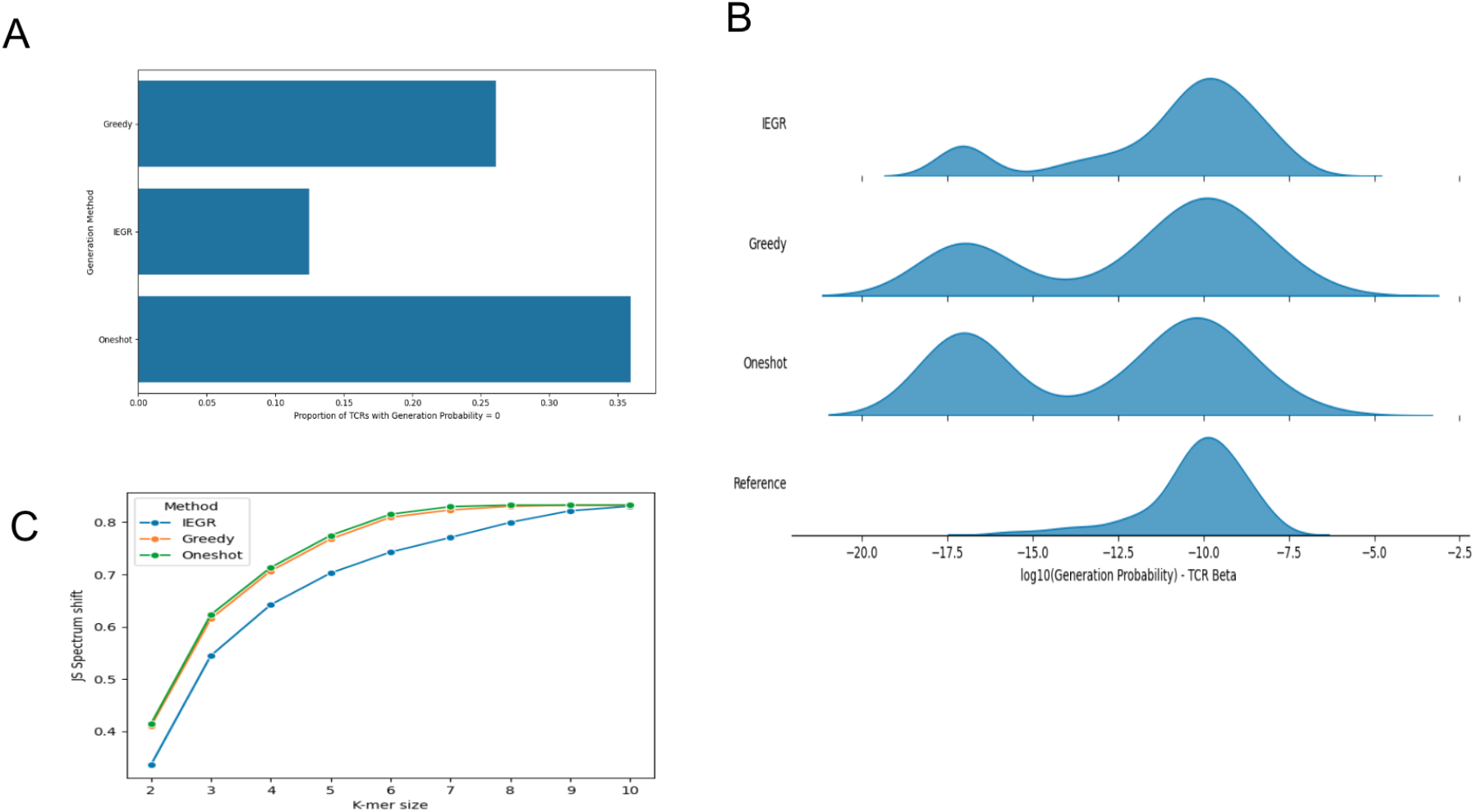
Sequence-level quality of generated TCRs. Comparison of generation methods targeting YLQPRTFLL/HLA-A*02:01. **(A)** Proportion of generated CDR3*β* sequences with zero V(D)J recombination probability (lower is better) obtained using the tool OLGA Sethna et al. (2019) **(B)** Distribution of log generation probability across methods; reference distribution shown for comparison. **(C)** K-mer spectrum shift (Jensen-Shannon divergence) relative to reference sequences across k-mer sizes (lower is better). IEGR produces sequences with generation statistics closest to natural TCR repertoires. K-mer spectrum shift calculated as outlined in Sarkar et al. (2024)

#### Experimental validation

We synthesized 30 designed TCR candidates (along with 6 controls), expressed them in TCR-knockout Jurkat cells, and measured dextramer binding by flow cytometry (Appendix F). IEGR produced one TCR with a staining above background (1/20, Design ID=30); other strategies yielded none. The IEGR designed TCR is novel (edit distance *≥*4 from training; Appendix E).

The experimental validation reflects the difficulty of jointly designing both CDR3*α* and CDR3*β*, unlike prior work that generates only one chain while fixing the other from known binders. Moreover, we note that ipTM scores do not reliably predict binding, underscoring that current in silico proxies lack resolution for true generative design benchmarking.

## 6. Discussion and Conclusion

We introduced a compositional pretraining framework demonstrating that protein language models *can* effectively model multi-component immune interfaces. Our two-stage curriculum provides a template for foundation models in domains with heterogeneous data availability.

### The prediction-generation gap

Our results reveal that strong zero-shot *prediction* does not trivially translate to reliable *generation*. The model discriminates binders from nonbinders substantially better than chance (0.76 AUROC) but is not yet capable of consistently generating strong binders. This asymmetry suggests that ranking requires capturing *necessary* conditions (sequence grammar, anchor constraints), while generation requires *sufficient* conditions demanding more paired data or thermodynamic calibration. We view this honest characterization as a contribution: it establishes realistic expectations and identifies where future effort should focus.

### Generalizable principles

Two principles extend beyond TCR-pMHC. First, compositional pre-training enables sample-efficient learning by projecting interaction learning onto a lower-dimensional manifold where limited paired supervision suffices. Second, confidence-based decoding respects biological hierarchy, naturally resolving conserved anchors before hypervariable positions. These principles may apply broadly to multi-component interfaces where paired data is scarce but marginal data is abundant.

### Limitations

Zero-shot epitope-specific design remains challenging. While all designed sequences are novel (edit distance *≥*4), we used existing binders as scaffolds; scaffold-free design is unsolved. Recognition performance varies across antigens, reflecting fundamental data sparsity (Lu et al., 2025). Recent estimates suggest 10^6^–10^8^ epitopes may be required for fully generalizable prediction (Delaunay et al., 2025); our approach may reduce but does not eliminate this data requirement.

### Conclusion

DecoderTCR demonstrates that compositional pre-training enables strong zero-shot *prediction* for TCR-pMHC interfaces, substantially outperforming generic protein language models. For *generation*, IEGR produces sequences passing structural and biological plausibility filters. Crucially, our experimental validation provides, to our knowledge, the first rigorous characterization of the prediction-generation gap, establishing a concrete benchmark for advancing generative immunology.

## Software and Data

Code for the DecoderTCR model will be made available at https://github.com/czbiohub-chi/DecoderTCR, and model checkpoints and other required files will be hosted at https://huggingface.co/biohub/DecoderTCR.

## Acknowledgements

We thank Kavita Kulkarni, Amanda Surya, Shaowen Zhu, Kyle Hippe, Tim Rand for helpful discussions. A.A.K. was supported in part by NIH DP2AI177884, a Chan Zuckerberg Investigator Award, and a Common Mechanisms of Autoimmunity Insight Award.

## Impact Statement

This work contributes to computational immunology with the goal of accelerating new therapies and diagnostics for cancer, autoimmunity, and infectious diseases. The ability to generate high-affinity TCRs *in silico* could reduce drug discovery time and cost, making personalized immunotherapies more accessible.

We acknowledge dual-use potential: generative capabilities for targeting cancer neoantigens could theoretically be misused to design receptors targeting healthy tissue or benign antigens. Additionally, training data is historically skewed toward MHC alleles common in European populations, risking inequitable distribution of therapeutic utility.

To mitigate these risks, we advocate for: (1) rigorous *in vitro* off-target screening against healthy tissue antigens before clinical application; (2) community prioritization of diverse immunogenomic datasets for equitable allele coverage; and (3) responsible model release under licenses restricting use to therapeutic research. We believe that with these safeguards, the therapeutic benefits of generative immunology outweigh the risks.

## A. Training Details

This appendix provides implementation details for the training procedures described in Section 3.2.

### A.1. Stage-Specific Masking Distributions

We use a masked residue prediction objective with a component- and stage-dependent masking distribution. For a sequence *x*, we sample a mask set *M⊆* {1, …, |*x*|} by independently masking position *i* with probability *p*(*i*) determined by its annotation (e.g., peptide, HLA pocket, CDR, or other). To take advantage of the structure of the interaction modality of the system of interest, we define:

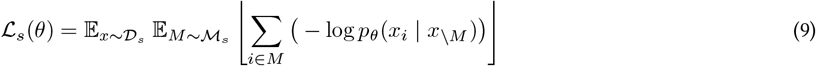

#### Stage 1 (Component specific)

Stage 1 uses two datasets: a peptide–MHC (pMHC) dataset *D*_pMHC_ and an unpaired TCR dataset *𝒟*_TCR_. For samples *x ∼ 𝒟*_pMHC_, we define the per-position masking probabilities as

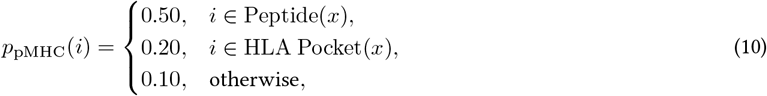

where Pocket(*x*) denotes the HLA binding pocket residues(exon2 & exon3). For samples *x ∼ 𝒟*_TCR_, we use

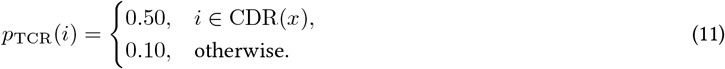

We sample *M* by masking each position *i* independently with the corresponding probability *p*_pMHC_(*i*) or *p*_TCR_(*i*), denoted *𝒫*_informed_.

#### Stage 2 (Cross-component interaction)

Stage 2 trains on a paired TCR–pMHC interaction dataset *𝒟*_paired_. For interaction samples *x ∼ 𝒟*_paired_, we define

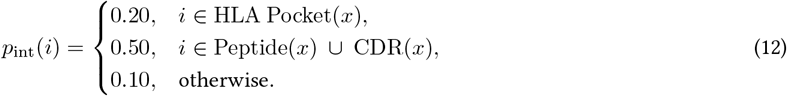

As in Stage 1, we sample *M* by independently masking each position *i* with probability *p*_int_(*i*).

### A.2. Optimization Configuration

We train with AdamW optimizer (learning rate 5 *×*10^*−*5^, *β*_1_ = 0.9, *β*_2_ = 0.999, weight decay 0.01). Batch size 4 per device across 96 H100 GPUs with BF16 mixed precision. Stage 1 processes 5.03*×* 10^11^ tokens. Stage 2 runs for 20 epochs. IEGR hyperparameters: *K* = 5*×*|*𝒰*_0_| refinement rounds, block size *b* = 3, temperature *τ* = 0.1. Hyperparameters selected on validation set (10% of paired data).

### A.3. Computational Cost

Model training processes around 5.03 *×* 10^11^ tokens required approximately 4,500 GPU hours on NVIDIA H100 Tensor Core GPUs, corresponding to roughly 1.6 *×* 10^22^ FLOPs (*≈* 1.6 *×* 10^10^ TFLOPs, or 16 exaFLOPs, BF16-equivalent).

## B. IEGR Algorithm Details

This section provides implementation details for Algorithm 1 in Section 3.4.

### B.1. Hyperparameters

### B.2. Implementation Details

#### Entropy computation

For masked positions *M*, predictive entropy quantifies uncertainty:

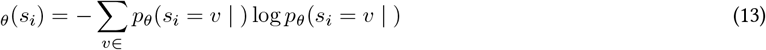

where *𝒱* is the 20-amino-acid vocabulary. We use natural logarithm, giving entropy in nats with maximum log(20) *≈* 3.0.

#### Temperature-scaled sampling

Given logits *𝓁*_*i*_ *∈* R^|*𝒱*|^ for position *i*, we sample:

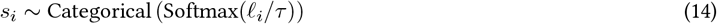

Lower *τ* concentrates probability mass on high-likelihood residues.

#### Block selection

In Phase 2 refinement, blocks are sampled uniformly at random from design positions *𝒰*_0_. Positions within a block are resampled with temperature *τ*

#### Motivation for Entropy-Guided Decoding Scheduling

Standard MCMC sampling on protein landscapes often suffers from slow mixing and kinetic trapping. Our IEGR schedule is a heuristic approximation inspired by confidence-based non-autoregressive decoding (Ghazvininejad et al., 2019). While we do not provide theoretical guarantees on mode coverage or ergodicity, the schedule is motivated by the biophysical hierarchy of protein folding: high-confidence residues (anchors) effectively constrain the conformational search space for lower-confidence loops. Empirically, we find this “confidence-first” filling reduces the likelihood of hallucinating invalid high-energy states early in the generation process compared to random-order Gibbs sampling.

## C. Data Preparation

This appendix details data collection and processing for the experimental setup described in Section 4.

### C.1. HLA Data Processing

HLA protein sequences were collected from the IPD-IMGT/HLA alignment database (Barker et al., 2023). Sequences were collected for 4 Class I genes (A, B, C, and E) and 6 Class II genes (DRA, DRB1, DPA1, DPB1, DQA1, DQB1). Eight-digit HLA allele names were simplified to four-digit allele resolution, then deduplicated based on unique protein sequences, keeping the longest sequence variant. A total of 17,065 HLA Class I alleles and 7,718 HLA Class II alleles were used for further processing.

### C.2. pMHC Data

#### C.2.1. Experimentally Validated

We collected experimentally validated pMHC pairs from Sarkizova et al. (2020); Tadros et al. (2023); Abelin et al. (2017); Pyke et al. (2021) for Class I HLAs and from The MHC Motif Atlas (Tadros et al., 2023) for Class II HLAs. Class I pairs were further filtered using MixMHCpred (binding score *−>* 2), and all samples were deduplicated based on peptide-allele pairs. We retained 451,828 unique pairs across 123 Class I HLA alleles and 548,858 unique pairs across 81 HLA Class II alleles.

#### C.2.2. Synthetic

For synthetic pMHC binders, human proteome sequences from the NCBI RefSeq database using the GRCh38.p14 assembly (Accession: GCF-000001405.40) were used to generate new peptides. The peptides of varying length (8–14 for Class I; 12–21 for Class II) were generated using a sliding window and scored with MixMHCpred across the 126 Class I alleles and MixMHC2pred across 75 Class II alleles. For Class I alleles, positive pairs were selected per allele and peptide length by retaining peptides with ≤1% rank and had top 1% scores with a minimum positive score threshold among all scored protein fragments, and negative pairs were randomly sampled from the bottom 50% of MixMHCpred scores per allele to match the distribution of positive pairs. Similarly, for Class II alleles, positive pairs were those ranked in the top 1% per allele and peptide length, and negative pairs were generated using weak-binding thresholds and those at the bottom 1% were retained. For Class II alleles, MixMHC2pred’s predicted peptide core was used, and samples were deduplicated based on the core. All the selected synthetic pairs were filtered to exclude any peptide-allele combinations present in the experimental datasets.

### C.3. TCR Data Processing

Each paired T cell receptor (TCR) alpha and beta chain was processed independently using Stitchr (Heather et al., 2022) and ANARCI (Dunbar and Deane, 2016). Only TCR pairs with complete V gene, CDR3 sequence, and J gene information available for both chains were retained for processing. For each chain, Stitchr generated the full-length TCR sequence from the input V gene, CDR3 sequence, and J gene calls. ANARCI was then used to annotate the CDR regions and validate the sequence generated by Stitchr. TCR pairs where either chain failed processing through Stitchr, or failed CDR3 regions validation by ANARCI, were excluded from further analysis. In datasets where full-length *α* and *β* chain sequences were already available (e.g., the OTS dataset), we performed an additional validation to confirm that sequences generated by Stitchr matched the original ones. For VDJdb, where we have the information on the full TCR-pMHC complex, we filtered out the samples with no available HLA allele for them.

### C.4. Unpaired TCR Repertoire Data

We collected and processed TCR sequences from Raybould et al. (2024) for the pre-training dataset. From the dataset, 1,511,895 distinct full-length TCR sequences were obtained after processing through Stitchr and ANARCI.

### C.5. TCR-pMHC Interaction Data

Experimentally validated immunogenic TCR-pMHC pair sequence data was collected from VDJdb (Bagaev et al., 2020), a curated database of T cell receptor sequences with experimentally validated antigen specificities (vdjdb.cdr3.net). We used the vdjdb-2024-11-27 release, downloaded from the official GitHub repository.

Similar to the pre-training dataset, the TCRs from VDJdb were processed through Stitchr and ANARCI. Only sequences that were successfully processed through both tools were retained. TCR entries with missing MHC allele information were excluded from the final dataset. Post-processing we obtained a total of 31,643 TCR sequences for further analysis.

### C.6. Quantitative pMHC binding data

Quantitative binding data were curated from the Immune Epitope Database (IEDB)(Vita et al., 2019) by filtering for peptides ligands to HLA-A, HLA-B, and HLA-C alleles with 9 amino acids in length and removing entries containing non-canonical residues. To stabilize variance across the broad range of binding affinities, *IC*_50_ values were transformed to a log_10_ scale.

### C.7. Dataset Summary

Table 5 summarizes the collected data across experimental and synthetic pMHC pairs for both Class I and Class II HLAs, as well as TCR datasets.

**Table 5.** Summary of experimental and synthetic dataset sizes.

| Dataset Type | Category | Count |
| --- | --- | --- |
| **Experimental pMHC (Class I)** | Total HLAs | 17,065 |
|  | HLAs with validated peptides | 123 |
|  | pMHC pairs | 451,828 |
| **Experimental pMHC (Class II)** | Total HLAs | 7,718 |
|  | HLAs with validated peptides | 81 |
|  | pMHC pairs | 548,858 |
| **Synthetic pMHC (Class I)** | Positive pairs | 9,464,613 |
|  | Negative pairs | 9,464,613 |
| **Synthetic pMHC (Class II)** | Positive pairs | 3,623,224 |
|  | Negative pairs | 20,432,290 |
| **TCR Datasets** | VDJdb | 31,643 |
|  | OTS | 1,511,895 |

**Table 6.**
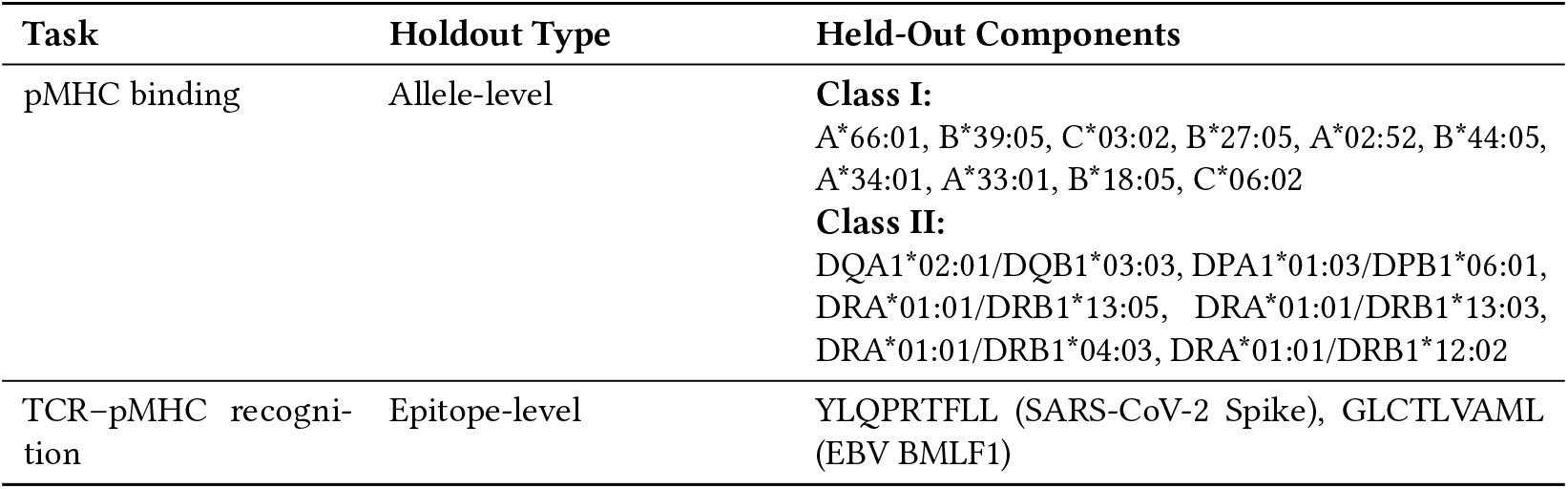
Summary of held-out data splits used for zero-shot evaluation.

**Table 7.** Masking strategy ablation on zero-shot pMHC binding. Mean *±* SD across held-out alleles.

| Model | Class I (10 alleles) |  | Class II (6 alleles) |  |
| --- | --- | --- | --- | --- |
|  | AUROC | AUPRC | AUROC | AUPRC |
| *Generic pLM* |  |  |  |  |
| ESM-2 (650M) | .41±.09 | .43±.05 | .51±.22 | .54±.17 |
| ESM-2 (3B) | .45±.11 | .46±.06 | .56±.18 | .56±.15 |
| *Uniform Masking (pMHC only)* |  |  |  |  |
| +pMHC (650M) | .97±.03 | .97±.03 | .87±.04 | .84±.05 |
| +pMHC (3B) | .98±.02 | .98±.02 | .90±.02 | .89±.03 |
| *Informed Masking (pMHC only)* |  |  |  |  |
| +pMHC (650M) | .98±.02 | .98±.03 | .88±.06 | .86±.05 |
| +pMHC (3B) | **.98±.03** | **.98±.03** | **.92±.02** | **.91±.03** |

**Table 8.** TCR Designs for YLQ Peptide with Updated Method Names.

| Design_ID | TRAV | TRBV | TRAJ | TRBJ | CDR3α | CDR3β | Edit-Dist | iPTM | Method |
| --- | --- | --- | --- | --- | --- | --- | --- | --- | --- |
| 30 | TRAV12-1 | TRBV7-8 | TRAJ30 | TRBJ2-2 | CVVNRDDKIIF | CASLDLNTGELFF | 4 | 0.79 | IEGR |
| 29 | TRAV12-1 | TRBV7-8 | TRAJ30 | TRBJ2-2 | CVVNRDDKIIF | CASLDGNTGELFF | 4 | 0.78 | IEGR |
| 15 | TRAV12-1 | TRBV7-9 | TRAJ34 | TRBJ2-2 | CVPYNTDKLIF | CASLEGNTGELFF | 5 | 0.76 | IEGR |
| 16 | TRAV12-1 | TRBV7-9 | TRAJ34 | TRBJ2-2 | CVPYNTDKLIF | CASLDSNTGELFF | 7 | 0.76 | IEGR |
| 17 | TRAV12-1 | TRBV7-9 | TRAJ34 | TRBJ2-2 | CVPYNTDKLIF | CASRLSNTGELFF | 7 | 0.76 | IEGR |
| 19 | TRAV12-1 | TRBV7-9 | TRAJ34 | TRBJ2-2 | CVPYNTDKLIF | CASLVSNTGELFF | 7 | 0.76 | IEGR |

**Table 8.**
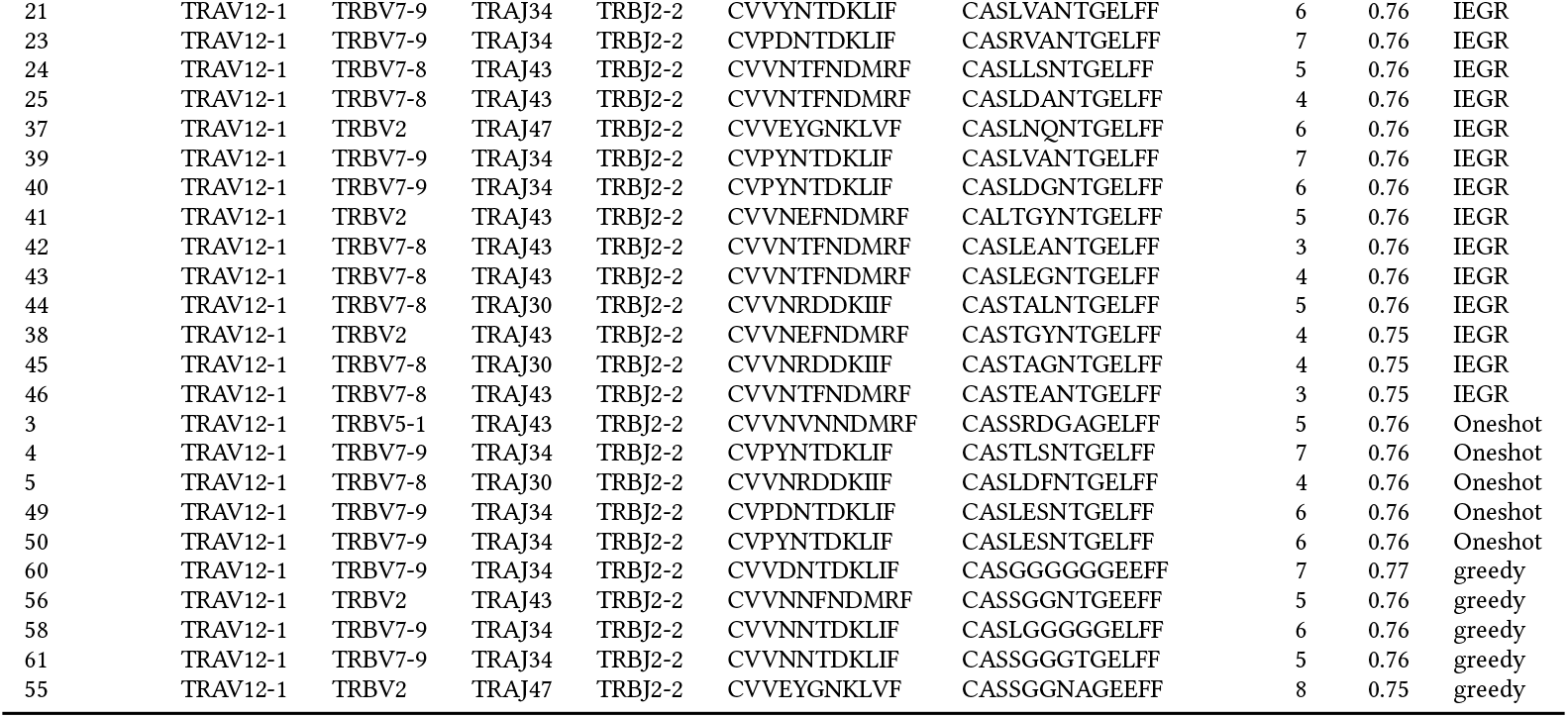
TCR Designs for YLQ Peptide with Updated Method Names.

### C.8. Held-Out Splits

To enable rigorous zero-shot evaluation, we constructed component-specific held-out datasets with no sequence overlap with the corresponding training data.

For the TCR-pMHC recognition task, we used a dataset generated by Messemaker et al. (2025) that validated a subset of VDJdb entries for the YLQPRTFLL and GLCTLVAML epitopes binding to HLA-A*02:01, providing experimentally confirmed binders and non-binders.

## D. Ablation Studies

This appendix provides additional ablation experiments supplementing the results in Section 5.

### D.1. Masking Strategy

To isolate the effect of informed masking, we trained on pMHC data alone (without TCR data) comparing uniform masking (15% of all positions) against informed masking (50% peptide, 20% HLA pocket, 10% elsewhere).

Informed masking provides consistent improvements, particularly for Class II alleles (+0.02 AUROC with 3B model). This confirms that concentrating supervision on binding-relevant positions provides useful inductive bias for learning MHC-specific binding preferences.

## E. Generated Sequences

This appendix provides the complete set of designed TCR sequences from the generation experiments in Section 5.5. We evaluate designed TCR sequences using both structural confidence metrics and sequence-level quality measures.

### E.1. Designed sequences selected for experimental validation

### E.2. Sequence Analysis

#### Design diversity and novelty

All validated binders are novel, with Levenshtein distance *≥* 4 from the nearest CDR3*αβ* recognizing YLQPRTFLL in VDJdb (mean 5.8 *±*1.4). Since all YLQPRTFLL-binding TCRs were held out during training, this confirms *de novo* design for an unseen epitope rather than memorizing known binders. Selected binders show mean pairwise distance of 4.2, indicating that IEGR explores diverse sequence solutions rather than converging to a single design motif.

## F. Experimental Protocols

This appendix details the wet-lab validation protocols for the experimental results reported in Section 5.5.

### F.1. Cell Lines

CD8^+^ TCR-knockout NFAT-Luciferase Reporter Jurkat cells (BPS Bioscience #78757), were engineered using CRISPR to knock out *FAS* creating a *TCR-/FAS*cells line. Cells were cultured in RPMI 1640 medium (Thermo Fisher Scientific 11875093) containing 10% fetal bovine serum and 1% penicillin-streptomycin and maintained in a 37°C incubator with 5% CO2.

### F.2. TCR Vector Design and Synthesis

Designed TCR sequences were individually synthesized and cloned by Twist Biosciences into their pTwist EF1 Alpha mammalian expression vector, with the TCR under control of the human EF1a promoter. Cloned plasmids were prepared by Twist Biosciences and used directly in nucleofection after resuspension to 500ng/uL in Tris-EDTA pH 8.0.

### F.3. TCR Expression/Nuclefection

*CD8+ TCR-/FAS*Jurkats were nucleofected using Lonza’s 4D nucleofector X-Unit in 16-well strip format using SE Cell Line 4D-Nucleofector® X Kit S (V4XC-1032) and pulse code CL-120. For each TCR-expressing plasmid, 200,000 cells each and 650ng TCR plasmid in 20 *µ*L SE combined + Supplement 1 (4.5:1), were pulsed, and after a 10 minute recovery in 80 *µ*L RPMI, plated in 400 *µ*L pre-warmed complete media. Cells were incubated as above for 48 hours, followed by dextramer staining and readout.

### F.4. Dextramer Staining and Flow Cytometry

Each full well of nucleofected cells was recovered after 48 hours and washed once with FACS Buffer (10% fetal bovine serum and 2mM EDTA in 1x phosphate buffered saline). Cells were then stained for 10 minutes at room temperature with 10 *µ*L of HLA-A*02:01/YLQPRTFLL-APC dextramer (Immudex WB05824). Cells were washed again, then stained with 1.2 *µ*L Brilliant Violet 421™ anti-human CD3 Antibody (Biolegend 300434) for 20 minutes at room temperature. Three more washes of 200 *µ*L were performed, followed by staining with 0.2 *µ*g/mL Propidium Iodide (Thermo Fisher Scientific P3566) for live/dead discrimination, and analysis on a SONY MA900 Cell Sorter. CD3+/APC+/PIevents exceeding non-binding control background were considered positive.

## G. Additional Results

This appendix provides extended results supplementing the main evaluation in Section 5.

